# Heterologous expression of the immune receptor *EFR* in *Medicago truncatula* reduces pathogenic infection, but not rhizobial symbiosis

**DOI:** 10.1101/171868

**Authors:** Sebastian Pfeilmeier, Jeoffrey George, Alice Morel, Sonali Roy, Matthew Smoker, Lena Stransfeld, J. Allan Downie, Nemo Peeters, Jacob G. Malone, Cyril Zipfel

## Abstract

Interfamily transfer of plant pattern recognition receptors (PRRs) represents a promising biotechnological approach to engineer broad-spectrum, and potentially durable, disease resistance in crops. It is however unclear whether new recognition specificities to given pathogen-associated molecular patterns (PAMPs) affect the interaction of the recipient plant with beneficial microbes. To test this in a direct reductionist approach, we transferred the *Brassicaceae*-specific PRR ELONGATION FACTOR-THERMO UNSTABLE RECEPTOR (EFR) from *Arabidopsis thaliana* to the legume *Medicago truncatula*, conferring recognition of the bacterial EF-Tu protein. Constitutive *EFR* expression led to EFR accumulation and activation of immune responses upon treatment with the EF-Tu-derived elf18 peptide in leaves and roots. The interaction of *M. truncatula* with the bacterial symbiont *Sinorhizobium meliloti* is characterized by the formation of root nodules that fix atmospheric nitrogen. Although nodule numbers were slightly reduced at an early stage of the infection in *EFR*-*Medicago* when compared to control lines, nodulation was similar in all lines at later stages. Furthermore, nodule colonization by rhizobia, and nitrogen fixation were not compromised by *EFR* expression. Importantly, the *M. truncatula* lines expressing *EFR* were substantially more resistant to the root bacterial pathogen *Ralstonia solanacearum*. Our data suggest that the transfer of EFR to *M. truncatula* does not impede root nodule symbiosis, but has a positive impact on disease resistance against a bacterial pathogen. In addition, our results indicate that *Rhizobium* can either avoid PAMP recognition during the infection process, or is able to actively suppress immune signaling.

**Significance Statement:** Crop engineering helps reducing the economic and environmental costs of plant disease. The genetic transfer of immune receptors across plant species is a promising biotechnological approach to increase disease resistance. Surface-localized pattern-recognition receptors (PRRs), which detect conserved characteristic microbial features, are functional in heterologous taxonomically-diverse plant species, and confer broad-spectrum disease resistance. It was unclear whether PRR transfer negatively impacts the association of the recipient plants with symbiotic microbes. Here, we show that a legume engineered with a novel PRR recognizing a conserved bacterial protein becomes more resistant to an important bacterial pathogen without significant impact on nitrogen-fixing symbiosis with rhizobia. This finding is of particular relevance as attempts to transfer this important symbiosis into non-legume plants are ongoing.

## Introduction

Plant pattern recognition receptors (PRRs) perceive conserved characteristic microbial features, termed pathogen- or microbe-associated molecular patterns (PAMPs or MAMPs), and trigger an immune response commonly referred to as PAMP- or pattern-triggered immunity (PTI). This confers basal disease resistance against adapted pathogens and plays a major role in non-host resistance against non-adapted pathogens (1, 2). Typically, plant PRRs are receptor kinases (RKs) or receptor-like proteins (RLPs), which consist of an extracellular ligand binding domain, a transmembrane domain, and an intracellular kinase (in the case of RKs) or a cytoplasmic C-terminal extension (in the case of RLPs) (1). While some PRRs, such as FLAGELLIN SENSING 2 (FLS2, which detects the PAMP epitope flg22 from bacterial flagellin) are present in all higher plant species, others have only evolved in certain plant families (1, 3). For example, the ELONGATION FACTOR-TU RECEPTOR (EFR), which recognizes the highly abundant and conserved bacterial protein EF-Tu (or the PAMP epitope elf18) and has been identified in *Arabidopsis thaliana*, seems to be present only in *Brassicaceae* (3). Plants recognise a wide variety of PAMPs, and it is becoming increasingly clear with the identification of new plant PRRs that many PRRs have evolved in a family- or even species-specific manner (1). Based on these observations, the ability to transfer novel PAMP recognition capabilities across plant species, families or even classes represents a promising biotechnological strategy to engineer broad-spectrum (and potentially durable) disease resistance in crops (1, 4–6). For example, the transgenic expression of *EFR* in other plant species, such as tomato (*Solanum lycopersicum*), *Nicotiana benthamiana*, wheat (*Triticum aestivum*), or rice (*Oryza sativa*) confers elf18 recognition and quantitative resistance to a range of bacterial pathogens including *Ralstonia solanacearum*, *Pseudomonas syringae*, *Xanthomonas perforans*, *X. oryzae* and *Acidovorax avenae* (7–11). In addition, the PRR XA21 (which recognises the tyrosine-sulfated peptide RaxX; (12)) from the wild rice *O. longistaminata* confers increased resistance against *Xanthomonas spp.* when expressed in banana (*Musa sp.*), sweet orange (*Citrus sinensis*) or *N. benthamiana* (13–15). Similarly, the PRR ELICITIN RECEPTOR (ELR) from the wild potato *S. microdontum* or the *A. thaliana* PRR RECEPTOR-LIKE PROTEIN 23 (RLP23, which recognises the taxonomically conserved peptide nlp20) confer increased resistance to the oomycete *Phytophthora infestans* when expressed in cultivated potato (*S. tuberosum*) (16, 17). These recent selected examples illustrate that PRRs normally restricted to specific plant taxonomic lineages remain functional when expressed in other plant species. Beyond the biotechnological usefulness of this property, this also illustrates that immune signaling components acting downstream of PRRs must be (at least partially) functionally conserved.

While plants must constantly defend themselves against potential invaders, they also form close interactions with beneficial microbes, in what is commonly referred to as the plant microbiome (18, 19). While all plants express PRRs as part of their innate immune system, it is however still unclear whether the engineering of novel PAMP recognition specificities through heterologous PRR expression affects the beneficial interaction of plants with commensal microbes.

The symbiosis between rhizobia and legumes is a defined and well understood interaction involving mutual communication. The symbiotic interaction starts with plant roots secreting chemical signals, including flavonoids, to attract host-compatible rhizobia. In turn, rhizobia produce symbiosis-inducing Nod factor, which is perceived by the plant and triggers two independent, yet coordinated, developmental processes: nodule organogenesis and bacterial infection (20, 21). Bacteria attach to the root hair tip and form micro-colonies, from which they invade the plant tissue by growing inside a tubular structure called an infection thread. In parallel to the root hair infection, plant cortical cells divide and develop a new organ, known as a nodule, which ultimately accommodates the rhizobia. In mature nodules, bacteria live as membrane encased bacteroids and fix atmospheric nitrogen making it available to the plant (20). The harmonious interplay between both organisms requires continuous signal exchange and can be terminated at various stages (22, 23).

Although the role of PAMP perception and immune signaling during symbiosis has not been extensively studied, there is accumulating evidence to suggest that rhizobia are initially perceived as potential invaders (23). The apparent overlap of components and concepts between immunity and symbiosis signaling pathways in legumes is both intriguing, and relevant to the question about the importance of PAMP recognition during these contrasting processes (24). Rhizobia are capable of eliciting PTI, because suspension cultures of *Mesorhizobium loti* can trigger in the legume *Lotus japonicus* defense-associated responses, such as ethylene production, MAP kinase activation and immune gene transcription in a similar way to the flg22 peptide (25). In addition, PTI signaling triggered by exogenous application of the *Pseudomonas aeruginosa*-derived flg22 peptide delays nodulation and reduces nodule numbers during the symbiosis between *L. japonicus* and *M. loti* (25). Transcriptional studies in different legume species also reported the transient upregulation of immune-related genes upon first encounter with its rhizobial symbiont, followed by a downregulation during the onset of symbiosis. For example, immune and stress-related genes in *M. truncatula* roots were upregulated 1 hour post-inoculation (hpi) and subsequently downregulated to a minimal level at 12 hpi in response to *S. meliloti* inoculation (26). Similarly, transcriptome analysis of root hair cells from soybean (27) and *M. truncatula* (28) showed induction of defense genes 24 hpi, then a marked reduction at later time-points after infection with *Bradyrhizobium japonicum* and *S. meliloti*, respectively. Interestingly, the early activation of plant defense may play a role in the selection of symbionts and endophytes versus pathogens during the early stages of the rhizobia-legumes interaction (29).

To test directly in a reductionist approach whether a novel PAMP recognition ability affects the symbiotic interaction between legumes and rhizobia, we expressed the *A. thaliana EFR* (*EFR*) gene in *M. truncatula* to engineer the perception of EF-Tu (or elf18 peptide) from its symbiont *S. meliloti*. After confirming EFR functionality, we then tested if *EFR* expression had an impact on rhizobial infection and the symbiotic interaction. While infection was unaffected, nodulation was slightly reduced at an early time-point, but recovered fully by the later stages of symbiosis. Importantly, rhizobia in nodulated *EFR*-*Medicago* plants fixed atmospheric nitrogen as efficiently as in control plants. Despite the lack of effect on rhizobial infection and nodulation, *EFR* expression conferred quantitative resistance to the bacterial root pathogen *R. solanacearum*, suggesting that the transfer of PRRs is an efficient biotechnological tool to confer increased legume resistance to pathogens, with minimal impact on symbiotic interactions.

## Results

### Transgenic expression of *EFR* in *Medicago truncatula* confers elf18 recognition in leaves and roots

To study the effect of *EFR* expression on symbiotic and pathogenic interactions with *M. truncatula*, we stably transformed *M. truncatula* ecotype R108 with p*CaMV35S*::*EFR-HA* by *Agrobacterium tumefaciens*-mediated transformation (30). Two independently transformed lines with a single insertion event were isolated (lines 26-8 and 18-1) and characterized alongside their respective null segregants as controls (lines 26-2 and 18-3). Transgenic *EFR*-Medicago plants showed similar growth and development as their control lines (Figure 1A), and EFR accumulation could be detected in leaf tissue by western blot analysis (Figure 1B). EFR specifically perceives the PAMP elf18 from various bacterial species (Figure S1), including the *M. truncatula* symbiont *S. meliloti*, and initiates immune signaling (7, 31). Transgenic *EFR*-*Medicago* plants responded to local treatment with the elf18 peptide by production of a transient burst of reactive oxygen species (ROS) in leaves (Figure 1C) and roots (Figure 1D). We also tested responsiveness to the PAMP flg22, and confirmed that all lines responded to the peptide (Figure S2), showing that the presence of EFR does not interfere with the function of an endogenous PRR (*i.e.* FLS2). These results show that EFR is functional in *M. truncatula* as it provides responsiveness to the PAMP elf18.

**Figure 1.**
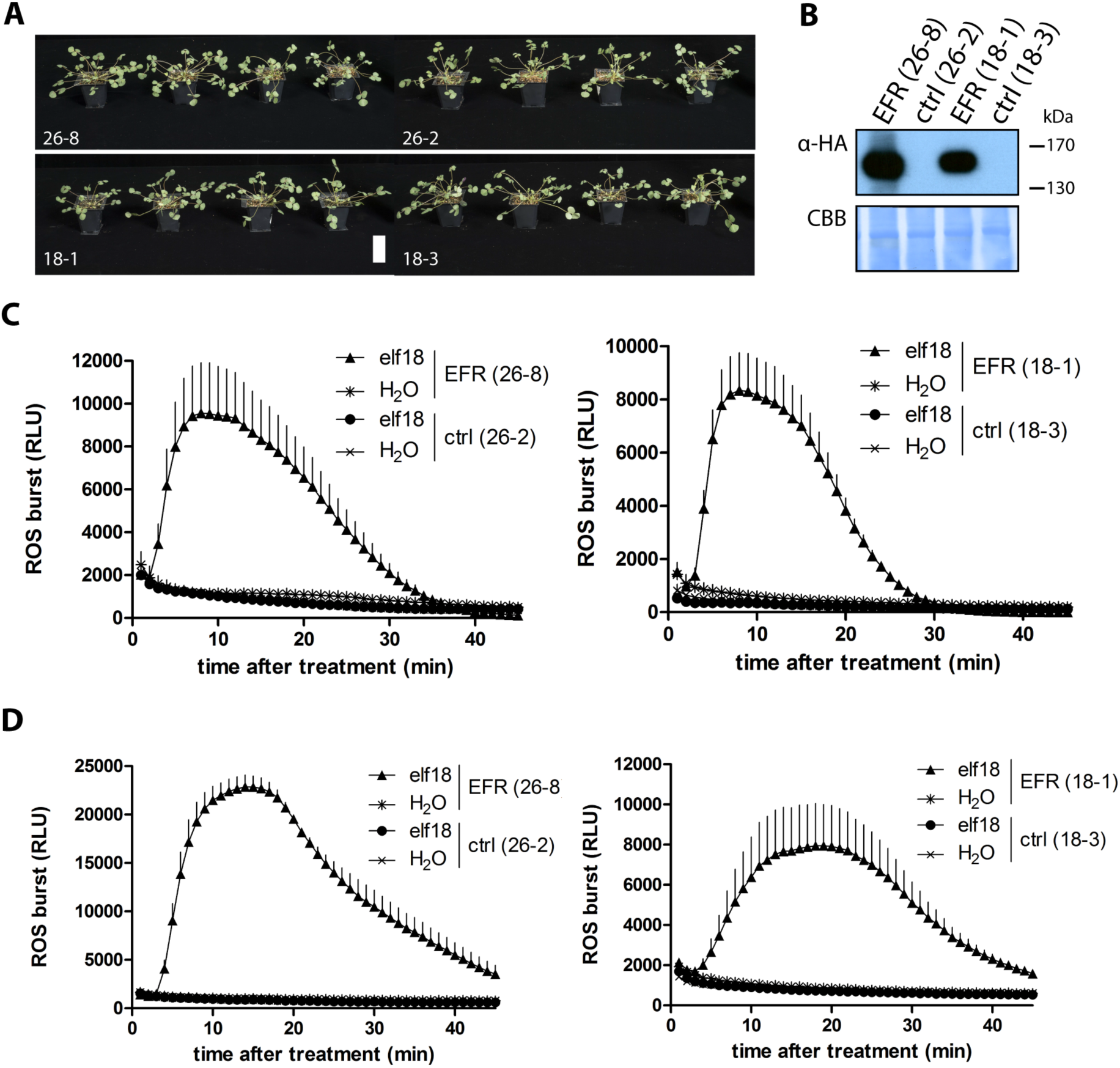
Transgenic *EFR*-*Medicago* responds to elf18 peptide. (A) Phenotype of two independent stable *EFR*-expressing *M. truncatula* lines, 26-8 and 18-1, and their null segregant control lines 26-2 and 18-3, respectively. White scale bar represents 5 cm. (B) Western blot of leaf material from *EFR*-*Medicago* lines (26-8 and 18-1) and control lines (26-2 and 18-3) using α-HA antibody to detect AtEFR-HA. Membrane was stained with Coomassie Brilliant Blue (CBB) as loading control. ROS burst was monitored in (C) leaf discs and (D) root segments from line 26-8 (left panels) and from line 18-1 (right panels) after application of 100 nM elf18 peptide and displayed as relative light units (RLU). Values are means ± standard error (*n*=8). The experiments were repeated twice.

### *EFR* expression does not affect the long-term symbiosis between *S. meliloti* and *M. truncatula*

We next tested whether heterologous expression of EFR affects the symbiosis between *S. meliloti* and *M. truncatula*. Expression of *EFR* in *M. truncatula* did not have a negative effect on plant growth after inoculation with *S. meliloti*, as the plant phenotype and fresh weight were similar in *EFR*-expressing and control plants when symbiosis was established at four weeks after infection (Figure 2A and B). *EFR* expression is driven by the ubiquitous *CaMV35S* promoter, and we were able to detect EFR accumulation in different root tissues, such as the main root, lateral roots and nodules (Figure S3). Next, we looked at different stages of the rhizobial infection and the nodulation process. Perception of PAMPs and PTI signaling presumably happens at the beginning of an infection, when the plant first encounters the microbe. We therefore tested whether EF-Tu recognition affects symbiotic interaction at this early stage. The formation of micro-colonies at the root hair tip, the number of infection threads and nodule primordia were similar between *EFR*-*Medicago* and the control lines (Figure 3A). Scoring total nodule numbers of the root at an early time-point (10 dpi) we observed a small, but significant, reduction in *EFR*-*Medicago* lines compared to control lines (Figure 3B). Total nodule numbers were reduced by 25% in line 18-1 and by 35% in line 26-8. Importantly, all nodules were colonized by rhizobia, as detected by staining for β-galactosidase activity in nodules colonized by the *S. meliloti* strain 1021-*lacZ*, and the spectrum of nodule morphology was similar for all lines. Notably, we observed a significant difference between the transformed null segregant control line 18-3 and the untransformed wild-type (Figure 3B), which encouraged us to use the null segregants as an appropriate control to avoid artefacts that could be linked to the genetic transformation and/or the associated *in vitro* culture process.

**Figure 2.**
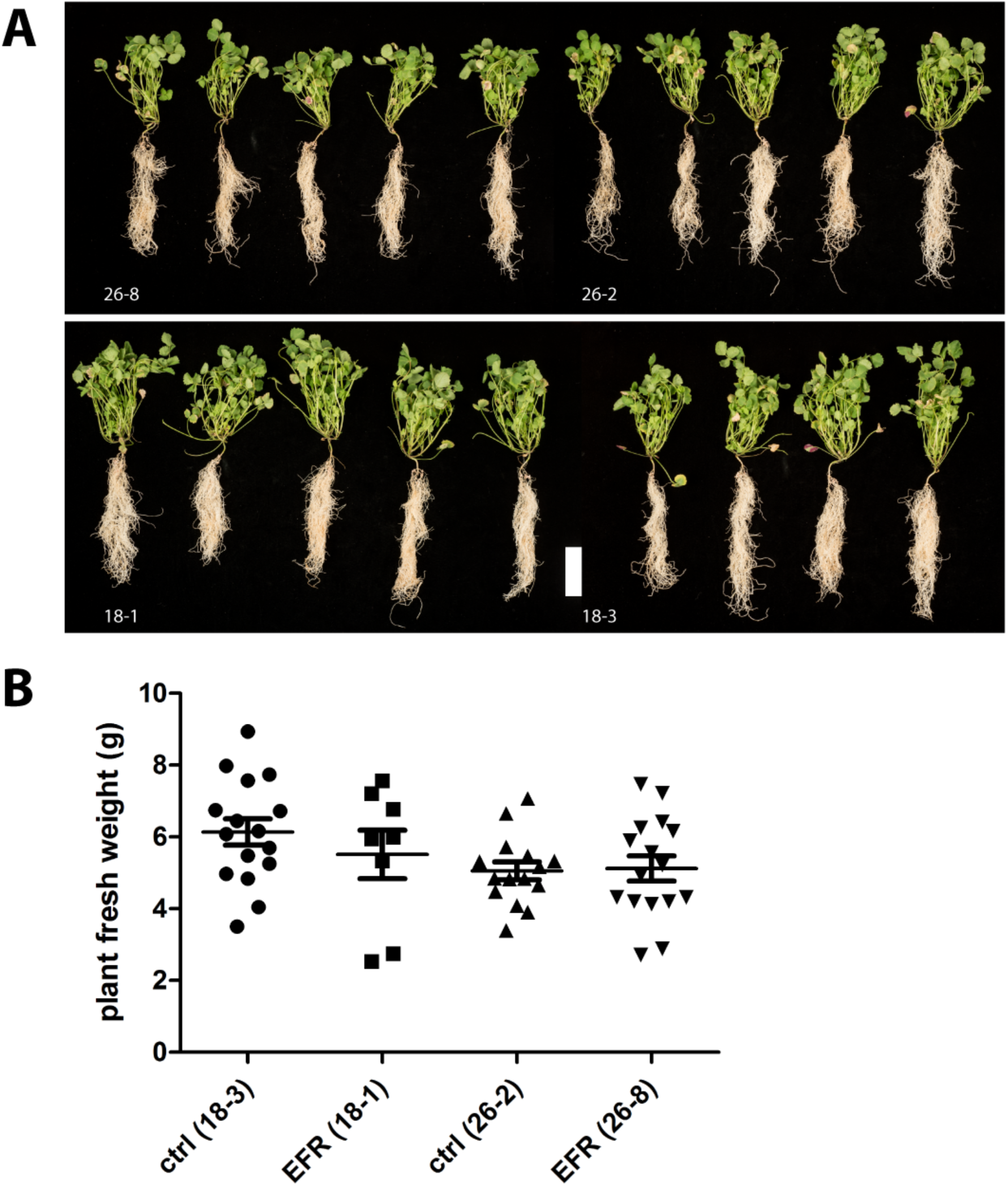
*EFR* expression does not affect development and fresh weight of *M. truncatula* infected with *S. meliloti*. (A) Plant pictures and (B) fresh weight was assessed of five-week old *M. truncatula* plants expressing *EFR* (26-8 and 18-1) and respective control lines (26-2 and 18-3) inoculated with *Sm*1021-*lacZ* and harvested at 28 dpi. White scale bar represents 5 cm. The experiments were repeated twice.

**Figure 3.**
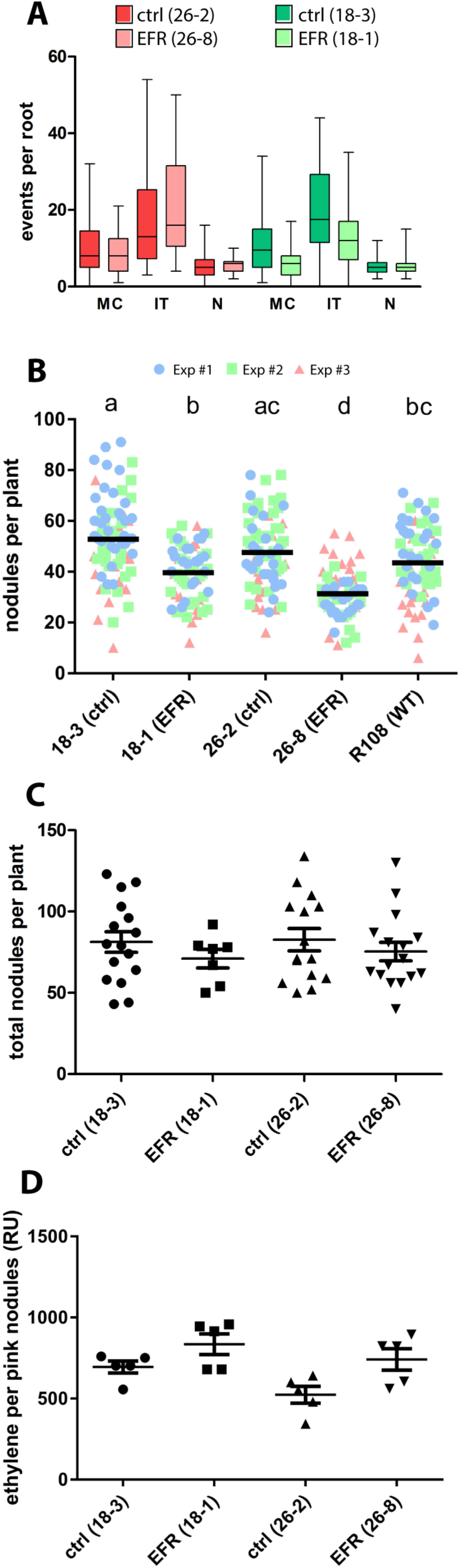
Symbiosis between *M. truncatula* and *S. meliloti* is not affected by *EFR* expression. (A) Infection events were scored at 7 dpi on roots of *M. truncatula* lines expressing *EFR* (26-8 and 18-1) and control lines (26-2 and 18-3) infected with *Sm*1021-*lacZ*. MC: micro-colonies. IT: infection threads. N: nodule primordia. Data from three independent experiments (each *n*=10) were combined. (B) Total nodules were scored at 10 dpi on roots of *M. truncatula* lines expressing *EFR* (26-8 and 18-1), control lines (26-2 and 18-3) and untransformed wild-type R108 infected with *Sm*1021*-lacZ*. Data from three independent experiments (each *n*=25) were combined. Letters indicate statistical significance groups with p<0.05 after One-way ANOVA (Kruskal-Wallis’s test and Dunn’s multiple comparison). (C) Total nodules were scored at 28 dpi on roots of *M. truncatula* lines expressing *EFR* (26-8 and 18-1) and control lines (26-2 and 18-3) infected with *Sm*1021-*lacZ*. One-way ANOVA with p<0.05 did not indicate statistical significant differences. (D) Acetylene reduction to ethylene was measured on whole plants of *M. truncatula* lines expressing *EFR* (26-8 and 18-1) and control lines (26-2 and 18-3) infected with *Sm*1021 *hemA::lacZ* at 28 dpi. Production of ethylene is displayed as relative units (RU) per pink nodules of each root system. One-way ANOVA with *p*<0.05 did not indicate statistical significant differences. The experiments were repeated twice.

We next assessed nodulation during the later stages of symbiosis. Four weeks after infection, symbiosis is well established and nodules are actively fixing atmospheric nitrogen (20). There was no difference in total nodule numbers between both *EFR*-*Medicago* lines and their respective controls (Figure 3C). In addition, we measured the enzymatic activity of rhizobial nitrogenase inside nodules, which can be used as an indicator of the nitrogen fixation rate (32). Notably, root systems from *EFR*-*Medicago* and from the control lines fixed nitrogen at similar rates (Figure 3D). In addition, the nodule morphology was similar in all lines, and no macroscopic signs of defense phenotypes or early senescence could be observed.

Together, our results thus indicate that *EFR* expression in *M. truncatula* may cause a slight initial delay in nodule formation, but overall does not negatively affect either rhizobial infection or long term-nitrogen-fixing symbiosis.

### *EFR* expression increases the resistance of *M. truncatula* to the bacterial root pathogen *R. solanacearum*

The proteobacterium *R. solanacearum* is a root pathogen that causes bacterial wilt disease in different plant species including *M. truncatula* (33, 34). *M. truncatula* plants infected with *R. solanacearum* develop disease symptoms such as chlorosis and wilting, ultimately leading to plant death (33). To test whether EFR can protect *M. truncatula* against bacterial pathogens, we infected *EFR*-*Medicago* and control lines with *R. solanacearum* GMI1000, and monitored disease progression and survival of the plants over several days. *EFR*-expressing plants displayed a consistently higher survival rate than plants from the control lines (Figure 4A and B). Although these results were only statistically significant for line 26-8 (based on Mantel-Cox test), we observed a tendency for higher disease resistance in line 18-1 across six independent experiments. A possible explanation for the enhanced disease resistance of line 26-8 compared to 18-3 may be the different EFR accumulation levels in these plants (Figures 1B and S3), which also translate into stronger elf18-induced ROS production in the root of line 26-8 compared to the line 18-1 (Figure S4).

**Figure 4.**
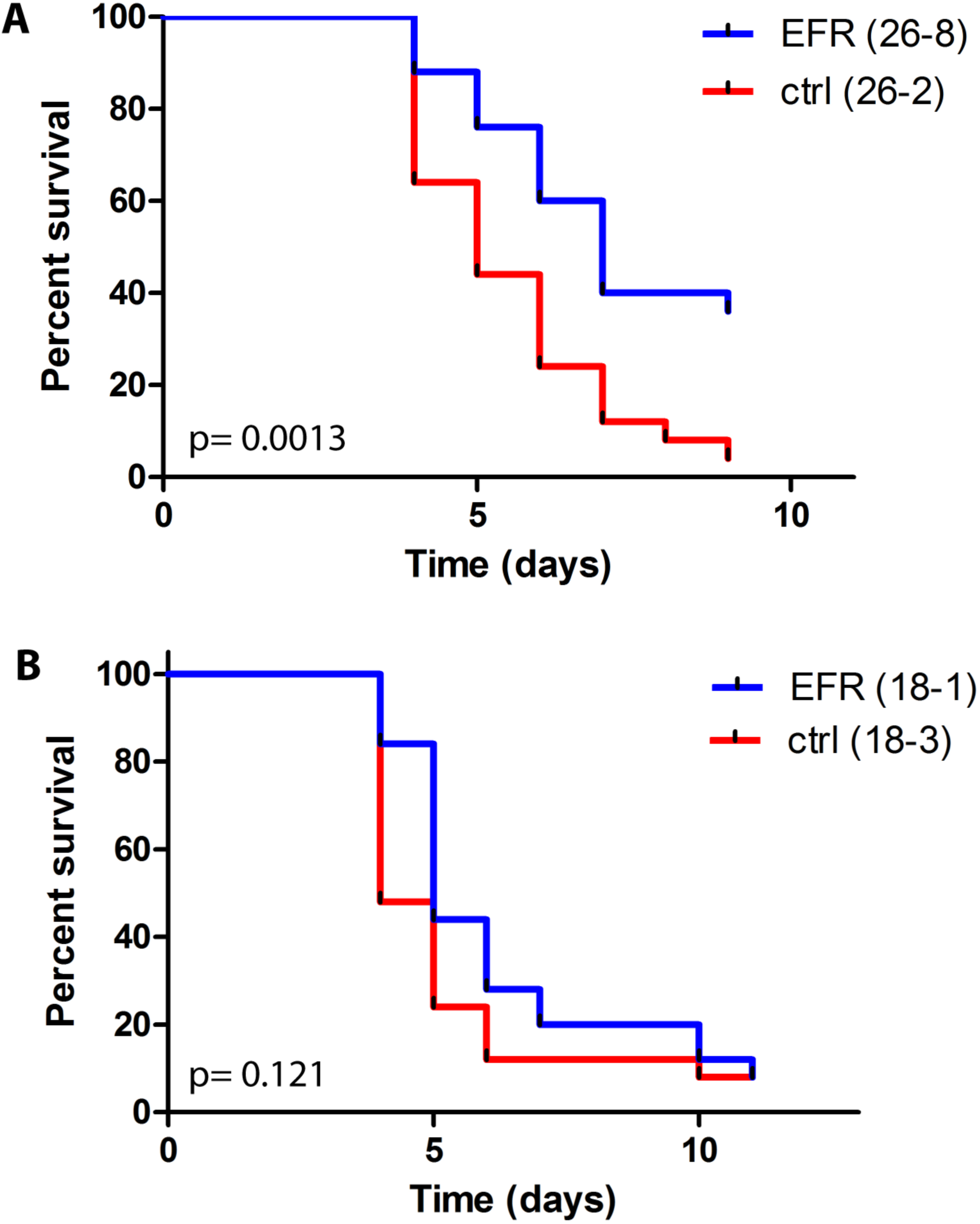
*EFR* expression in *M. truncatula* provides quantitative resistance against the pathogen *R. solanacearum*. (A) *M. truncatula* lines expressing *EFR* 26-8 and control line 26-2 were infected with *R. solanacearum* GMI1000 and disease symptoms assessed daily. Survival rate is displayed over 9 days and statistical analysis performed with Mantel-Cox test, p=0.0013 (*n*=25). Experiment was repeated four times with similar results. (B) *M. truncatula* lines expressing *EFR* 18-1 and control line 18-3 were infected with *R. solanacearum* GMI1000 and disease symptoms assessed daily. Survival rate is displayed over time and statistical analysis performed with Mantel-Cox test, *p*=0.121 (*n*=25). Experiment was repeated six times with similar tendency.

Overall, as previously observed with other *EFR*-expressing plant species infected with bacterial pathogens (7–11), EFR conferred increased quantitative resistance, delayed disease progression and increased survival under our experimental conditions.

## Discussion

Many PRRs have been successfully expressed in heterologous hosts across taxonomically diverse plant species to improve disease resistance, and PRRs have thus become attractive tools as part of the biotechnological arsenal to genetically engineer disease resistance in crops (1, 4–6). It was however still unclear whether such PRR transfer may negatively impact the association of the recipient plants with symbiotic microbes.

In this study, we generated transgenic *M. truncatula* lines that express the heterologous PRR *EFR* to confer recognition of the endogenous PAMP EF-Tu from the symbiotic bacterium *S. meliloti*. These *EFR*-*Medicago* plants recognized elf18 peptide and initiated a PAMP-induced ROS burst in both leaves and roots (Figure 1C and D). As previously reported (8, 15), these results indicate components involved in the PTI signaling pathway (and in PRR biogenesis) are also present and conserved in the root and leaves of *M. truncatula*, as they enable the functionality of EFR.

Legumes benefit from the symbiosis with rhizobia during growth in nitrogen-limited soil due to the additional bioavailable nitrogen supplied by the bacteria. The growth and development of plants with an established symbiosis was similar between transgenic *EFR*-*Medicago* and control lines. Therefore, we concluded that *EFR* expression did not exert a negative effect on growth of infected plants, or on long-term symbiosis (Figure 2A and B). Likewise, early interaction events were unaltered by *EFR* expression. There was no major difference in the occurrence of micro-colonies, infection threads and nodule primordia formation between *EFR*-*Medicago* and control lines (Figure 3A). Although infection and nodulation are triggered simultaneously, both processes belong to different developmental programs (35, 36). Interestingly, at an early time-point after rhizobial infection nodule numbers were slightly reduced in *EFR* expressing lines (Figure 3B). However, at later stages of symbiosis nodule numbers were similar in all lines (Figure 3C). These data indicate that while nodulation might be delayed at early stages, the expression of *EFR* did not impede long-term nodulation. Importantly, the nitrogen-fixation capacity of nodules was unaffected by *EFR* expression (Figure 3D).

The absence of detrimental effects of *EFR* expression on rhizobial nitrogen-fixing symbiosis may at first appear counterintuitive. Indeed, the EF-Tu-derived EFR ligand elf18 from *S. meliloti* is able to induce immune responses (7, 31), and it has previously been shown that elicitation of PTI using exogenous PAMP treatment can affect the interaction between rhizobia and legumes. For example, application of flg22 peptide or *M. loti* cells to *L. japonicus* triggered similar PTI responses leading to a delay in nodule formation and reduced nodule numbers (25). Furthermore, co-inoculation experiments in *M. truncatula* recently showed that the pathogenic bacterium *P.* syringae pv. *tomato* (*Pto*) DC3000 induces immune responses and suppresses the establishment of the symbiosis with *S. meliloti* (37). In addition, constitutive activation of immune responses in *M. truncatula* in specific mutants or over-expression lines impairs nodule formation and symbiosis (38–43). Interestingly, nodulation in *M. truncatula* was only impaired when plants were co-treated with the PAMP peptide flg22 together with *M. loti*, but not when flg22 was applied after symbiosis was established. PAMP treatment seems to delay nodule development rather than impairing rhizobial fitness, as spontaneously nodulating *snf1* mutant plants were similarly affected by flg22 treatment (25). It thus appears that the timing of PTI activation and symbiotic signaling may be important to study the impact of PAMP recognition on symbiosis. In this context, it is important to note that *EFR* expression in recipient transgenic plants does not seem associated with constitutive activation of immune responses, as no detrimental effects on plant growth or development have ever been observed in these plants in either axenic or non-sterile soil conditions (7–9, 11) (Figures 1A and 2). Thus, our findings that *EFR* expression negatively affects early nodulation but not infection events or nodule numbers, as well as the observation that nitrogen fixation was unchanged at later stages of symbiosis, support the notion that the perception of rhizobial PAMPs might have an early transient effect on plant nodulation but does not compromise rhizobial fitness, infection, or the ultimate establishment of a functional nitrogen-fixing symbiosis during the natural infection process.

Our data therefore suggest that during a natural infection process, *S. meliloti* either evades EF-Tu recognition or actively suppresses PTI in the host. While rhizobia are known to carry a flg22 allele of the flagellin gene that is not recognized by the plant FLS2 receptor (44), the EF-Tu-derived elf18 peptide from *S. meliloti* is recognised by EFR (7, 31), suggesting that an immune evasion strategy is not conceivable here. Despite the manifold evidence that *EFR* expression provides efficient disease resistance, it is still unclear how the intracellular EF-Tu protein (and by extension the elf18 epitope) gets exposed to the EFR receptor during infection (10). EF-Tu has however been found in the secretome of different bacterial species (45, 46), and an active role for EF-Tu has been suggested during effector translocation via the type-6 secretion system (T6SS) in *P. aeruginosa* (47). Interestingly, EF-Tu was recently identified in bacterial outer membrane vesicles, which was linked to the ability of these vesicles to induced EFR-dependent immune responses in *A. thaliana* (48), illustrating a possible mechanism by which this potent PAMP can be released. We cannot therefore completely exclude the possibility that rhizobia have evolved strategies to control EF-Tu secretion.

Previous transcriptomic studies indicate that rhizobia initially elicit an immune response, which is then suppressed as symbiosis proceeds (26–28). In addition, co-inoculation with *S. meliloti* suppresses immune responses normally triggered by *Pto* DC3000 in *M. truncatula* (37). These results suggest that rhizobia have active mechanisms to suppress PTI. Plant pathogenic bacteria can suppress host immunity by secretion of effectors, many of which interfere with the canonical PTI pathway at different stages (49). Many of these PTI-suppressing effectors are translocated within plant cells via the type-3 secretion system (T3SS) (50). While there is evidence suggesting that rhizobia also use effectors to suppress plant immunity, only a few rhizobial effectors (Nop proteins) have been characterized (51). Rhizobial genomes encode several different secretion pathways, and the importance of T3SS, type-4 secretion systems (T4SS) and T6SS for symbiosis has been demonstrated genetically in certain rhizobial species (52–54). *Sinorhizobium* sp. NGR234 translocates multiple type-3 secreted effectors including NopM, NopL, NopP and NopT to interfere with immune signaling (55–58). However, the *S. meliloti* strain 1021 used in our study, *Sm*1021, only carries a T4SS gene cluster for translocation of effectors into host cytoplasm, and deletion mutant studies showed no impact of mutating the T4SS on nodulation (59, 60).

Other bacterial mechanisms have been suggested to suppress PTI (23). For example, Nod factors not only play a role in initiating and maintaining symbiosis signaling, but also in suppression of plant immunity (61). Application of *B. japonicum* Nod factor to the non-host plant *A. thaliana* resulted in reduced accumulation of the immune receptors FLS2 and EFR at the plasma membrane (61). Although this partial suppression of PTI seems to be conserved in legume and non-legume plants, the impact on rhizobial-legume root infection has not been directly tested. Contrary to these findings, we actually detected EFR accumulation in the nodules of our transgenic *M. truncatula* plants (Figure S3). Exopolysaccharide (EPS) production is a common factor among plant-associated bacteria and has been previously associated with evasion of PTI (62, 63). Cell surface polysaccharides such as EPS, lipopolysaccharides (LPS) and cyclic β-glucans have been implicated in facilitating symbiotic interaction (64–66). Notably, the EPS receptor EPR3 from *L. japonicus* specifically detects EPS from its symbiont and acts as a positive regulator of infection (67, 68). Mutant strains defective in cell surface polysaccharides result in impaired infections or ineffective nitrogen-fixing nodules (62). For example, the succinoglycan-deficient *Sm*1021 *exoY* mutant induces immune-related genes more strongly than wild-type, indicating a possible involvement of succinoglycan in the suppression of immunity (69). Additionally, purified LPS from *S. meliloti* can suppress ROS burst in *M. truncatula* suspension cells treated with invertase (64). However, the phenotypes of cell surface polysaccharide mutants are often difficult to interpret, because they seem to be specific for the type of polysaccharides, the rhizobial species and the host plant. It is therefore likely that cell surface polysaccharides contribute to the symbiotic interaction in multiple ways in addition to facilitating immune evasion (70). It will be thus interesting to investigate in future studies the exact mechanisms employed by rhizobia to suppress PTI. It is also conceivable that legumes themselves specifically suppress PTI in a local and timely manner in response to rhizobial signals to facilitate infection by their symbionts. For example, limiting trafficking of PRRs into the infection thread and/or peribacteroid membranes could restrict recognition of those PAMPs.

The transfer of EFR has already been shown to confer increased quantitative resistance against different bacterial pathogens in a wide range of plant species, including tomato, *N. benthamiana*, wheat and rice (7–11). The present study expands this list and reveals that *EFR* is also functional when expressed in legumes (at least as demonstrated here for *M. truncatula*) and increases the survival of *M. truncatula* plants upon infection by the root bacterial pathogen *R. solanacearum* (Figure 4A,B). Thus, together, our data demonstrate that *EFR* expression can protect *M. truncatula* from the destructive root pathogen *R. solanacearum*, without compromising the overall symbiotic interaction with *S. meliloti*, which allows fixation of atmospheric nitrogen. Our results suggest that legumes can be engineered with novel PRRs without affecting the nitrogen-fixing symbiosis, and may also be relevant in the future as attempts to transfer this important symbiosis into non-legume plants are currently ongoing (Zipfel & Oldroyd, 2017). More generally, it also illustrates that the transfer of PRRs across plant species do not necessarily come at a cost for the plant, but actually increases its fitness when faced with aggressive pathogens. It will be interesting in the future to expand the reductionist approach used in this study to test whether heterologous PRR expression affects the composition and function of other commensals in the plant microbiome. A potential effect on the microbiome would then however need to be reconciled with the absence of obvious growth defects of transgenic plants expressing PRRs in non-sterile soil, and counterbalanced in an agricultural sense against the benefit conferred by PRR transfer in term of disease resistance under strong pathogen pressure.

## Material and Methods

### Bacterial growth

*S. meliloti* strain 1021 (also known as *Rm*1021) *pXLGD4* p*hemA::lacZ* (*Sm*1021-*lacZ*) (71) was grown at 30°C in TY medium (tryptone 5 g/L, yeast extract 3 g/L) containing appropriate antibiotics, streptomycin 50 μg/mL and tetracycline 12.5 μg/mL. *Ralstonia solanacearum* strain GMI1000 was grown at 28°C in complete BG medium (bacto peptone 10 g/L, casamino acid 1 g/L, yeast extract 1 g/L).

### Plant growth

*M. truncatula* seeds were scarified with 98% sulfuric acid treatment for 8 min, extensively washed with water and then surface-sterilized with 10% NaOCl for 2 min. After washing with sterile water, the seeds were left for 3 hours to imbibe water before being placed on inverted agar plates in the dark for 3 days at 4°C and subsequently germinated over-night at 20°C. For sterile growth, seedlings were transferred to squared 1% agar plates and sandwiched between two Whatman filter papers (GE Healthcare, UK). Plates were incubated vertically in a growth chamber at 21°C, with a 16h light period and 80% rel. humidity.

For growth in soil, germinated seedlings were transferred to sterile 1:1 mixture of terragreen (Oil-dry UK ltd, UK) and sharp sand (BB Minerals, UK) for rhizobial infections, in loam based compost (John Innes Cereal Mix) for seed bulking, or in Jiffy Peat Pellets for inoculation with *Ralstonia solanacearum*. Plants were grown in controlled environment chambers with a 16-hour photoperiod at 20°C with 120-180 μmol m^−2^ s^−2^ light intensity and 80% rel. humidity.

### Stable transformation of *Medicago truncatula*

*M. truncatula* ecotype R108 was stably transformed by *A. tumefaciens* AGL1 carrying the recombinant binary vector *pBIN19*-*CaMV35S*::*EFR-HA* (7). Plant transformation was carried out as previously described (30) with some minor, but important, changes; *in vitro* grown plants, only, were used for the transformations, excised leaves were sliced through with a scalpel and not vacuum infiltrated, the *A. tumefaciens* culture was used at OD_600_=0.4 and re-suspended in SH3a broth with 300 μM acetosyringone, the leaflet explants were submerged in the bacterial suspension for 20 min only, shaking in the dark, leaves were co-cultivated and callus generated with their adaxial surface in contact with the media, explants were washed in SH3a media broth post co-cultivation, excess *Agrobacteria* was eliminated on solid media using 320 mg/L ticarcillin disodium/potassium clavulanate and finally, callus growth was carried out in the dark for 8 weeks rather than 6. Five transgenic plants were recovered by somatic embryogenesis, rescued and selected on kanamycin plates. Homozygous plants were identified by quantitative real-time PCR of a segregating T1 population and confirmed in the T2 stage by responsiveness to elf18 peptide. Two homozygous lines with only a single insertion locus carrying two *EFR* copies were identified and used for physiological and symbiotic characterizations. In addition, null segregants were isolated for each primary transformant and used as control lines. All experiments were performed with plants from the T3 population.

### Rhizobial infections

Plants were grown in pots (50 or 80 mL volume) in terragreen/sharp sand mix for 2 days (infection thread counting) or 7 days (nodule counting and acetylene reduction measurements) before inoculation with *Sm*1021-*lacZ*. Bacteria were grown in TY to OD_600_=1.5, washed in 10 mM MgCl2 and diluted to OD_600_=0.0002. Plants were inoculated with 5 mL of *S. meliloti* suspension equally spread across the pot. Plants were harvested at indicated time-points, carefully rinsed with water and stained with X-Gal for visualization of LacZ activity. Stained nodules were counted under a stereo microscope. For late-time point experiments (*e.g.* 28 dpi), plants were grown in bigger pots (500 mL volume) to allow enough space for root development and were inoculated with 10 mL *S. meliloti* diluted to OD_600_=0.0002. Plant nodules were scored visually and were not stained.

### X-Gal staining of infection structures and nodules

For staining of infection threads on plants grown in terragreen/sand mixture, whole roots were detached from shoot and placed in fixing solution containing phosphate buffer, pH=7 (61 mM Na_2_HPO_4_, 39 mM NaH_2_PO_4_, 10 mM KCl, 1 mM MgCl2) and 2.5% glutaraldehyde. Vacuum was applied for 5 min and roots were incubated for 1 h at room temperature. Roots were washed three times in phosphate buffer before staining solution (5 mM K_4_[Fe(CN)_6_], 5 mM K_3_[Fe(CN)_6_], 0.08 % X-Gal (Formedium, King’s Lynn, UK) in phosphate buffer) was added and roots incubated in the dark at 30°C over-night. Roots were washed in phosphate buffer three times and stored at 4°C until analysis.

Stained infection structures were scored in brightfield mode using a Leica DMR microscope (Wetzlar, Germany). The infection events were classified into three categories: micro-colony formation at curled root hair, elongated and ramified infection threads and nodule primordia.

### *Ralstonia solanacearum* infection of *Medicago truncatula*

After germination, *M. truncatula* plants were transferred and grown in jiffy peat pellets. For inoculation, the 1/3 bottom half of the Jiffy pots was severed then soaked in a *R. solanacearum* solution at OD_600_=0.1. Potting soil was used to absorb the remaining inoculum and spread on the bottom of the tray before putting Jiffy pots back on. Disease symptoms were scored daily. Statistical analysis was performed as previously outlined (72).

### Acetylene Reduction Measurements

Nitrogenase activity was determined by gas chromatography measuring the enzymatic conversion of acetylene gas to ethylene as previously described (73). Infected plants at 28 dpi, placed in a 50-mL plastic vial and sealed with a rubber lid. Acetylene gas (BOC, Manchester, UK) was injected into the vials with 2% (v/v) final concentration, incubated for 1 h at 23°C and 1 mL sample taken for analysis. Conversion of acetylene to ethylene by rhizobial nitrogenase was recorded on a Clarus 480 (Perkin Elmer) gas chromatograph with N_2_ as the carrier gas set to a flow rate of 25 mL min^−1^, a HayeSep N 80/100 mesh column, connected to a flame ionisation detector at 100°C. Acetylene was applied in excess and peak areas of ethylene were quantified using TotalChrom Workstation software (Perkin Elmer) and displayed as relative units.

### ROS burst measurements

*M. truncatula* seedlings were grown sterile for 7 days under 16-hour photoperiod at 21°C. Roots were cut into 3 mm segments and recovered in water over-night. Alternatively, leaf discs were sampled from soil-grown 4–5 weeks-old plants and recovered in water. ROS burst was measured as described previously. The water was replaced with solution containing 200 μg/mL horseradish peroxidase (HRP) (Sigma-Aldrich) and 1 μM L-012 (Sigma-Aldrich), incubated for 5 min and topped up with solution containing flg22 or elf18 peptide (EZBiolab, Westfield, IN, USA) with 100 nM final concentration. Luminescence was recorded over 45 min using a charge-coupled device camera (Photek Ltd., St Leonards on Sea, East Sussex, UK).

### Immunoblot analysis

Plant tissue was ground in liquid nitrogen and protein was extracted using a buffer containing 100 mM Tris-HCl, pH 7.2, 150 mM NaCl, 5 mM EDTA, 5% SDS, 2 M urea, 10 mM DTT and 1% (v/v) Protease Inhibitor Cocktail (P9599, Sigma-Aldrich), boiled for 10 min and debris removed by centrifugation for 2 min at 17.000 rpm. Protein samples were separated on an 8% sodium dodecylsulfate polyacrylamide gel electrophoresis (SDS-PAGE) and blotted on polyvinylidene difluoride (PVDF) membrane (Thermo Fisher Scientific). Immunoblotting was performed with α-HA-horseradish peroxidase (HRP) antibody (3F10, Roche) diluted 1:2000 in 5% milk in TBS with 0.1% (v/v) Tween-20. Blots were developed with Pierce ECL pico Western Blotting substrate (Thermo Fisher Scientific). Equal loading of protein was determined by Coomassie Brilliant Blue staining of the blotted membrane.

## Acknowledgements

The authors thank technical assistance from the John Innes Centre Horticultural Services, and helpful discussions with members of the Zipfel and Malone laboratories. SP is funded by a studentship from the Norwich Research Park. Research in the Malone and Zipfel laboratories is supported by BBSRC Institute Strategic Program Grant BB/J004553/1. The Zipfel laboratory is further supported by the Gatsby Charitable Foundation. The work done at LIPM, France was supported by the Laboratoire ďExcellence (LABEX) TULIP (ANR-10-LABX-41). JAD thanks the John Innes Foundation for an Emeritus fellowship.

